# The reliability of the isotropic fractionator method for counting total cells and neurons

**DOI:** 10.1101/627869

**Authors:** Kleber Neves, Daniel Menezes, Danielle Rayêe, Bruna Valério-Gomes, Pamela Meneses Iack, Roberto Lent, Bruno Mota

**Author notes:** These authors contributed equally to this work (Kleber Neves, Daniel Menezes, Danielle Rayêe, Bruna Valério-Gomes).

## Abstract

**Background:** The Isotropic Fractionator (IF) is a method used to determine the cellular composition of nervous tissue. It has been mostly applied to assess variation across species, where differences are expected to be large enough not to be masked by methodological error. However, understanding the sources of variation in the method is important if the goal is to detect smaller differences, for example, in same-species comparisons. Comparisons between different mice strains suggest that the IF is consistent enough to detected these differences. Nevertheless, the internal validity of the method has not yet been examined directly.

**Method:** In this study, we evaluate the reliability of the IF method for the determination of cellular and neuronal numbers. We performed repeated cell counts of the same material by different experimenters to quantify different sources of variation.

**Results:** In total cell counts, we observed that for the cerebral cortex most of the variance was at the counter level. For the cerebellum, most of the variance is attributed to the sample itself. As for neurons, random error along with the immunological staining correspond to most of the variation, both in the cerebral cortex and in the cerebellum. Test-retest reliability coefficients were relatively high, especially for cell counts.

**Conclusions:** Although biases between counters and random variation in staining could be problematic when aggregating data from different sources, we offer practical suggestions to improve the reliability of the method. While small, this study is a most needed step towards more precise measurement of the brain’s cellular composition.

**Highlights:**

- Most variance in cell counts was between counters (η = 0.58) for cerebral cortices.
- For cerebella, most of the variance was attributed to the samples (η = 0.49).
- Variance in immunocytochemical counts was mostly residual/random (η > 0.8).
- Test-retest reliability was high (same counter, same sample).
- Practical suggestions are offered to improve the reliability of the method.

## 1 Introduction

Much of a biological system can be explained by its architecture – from the ultrastructure of its cells to how the cells are organized within the parenchyma. The central nervous system (CNS) has an extremely complex cytoarchitecture, composed mainly of two cell types, neuronal and glial cells, and dozens of cellular subsets, each one performing different functions within the circuits.

When studying such a complex system, different hierarchical levels may be explored in order to provide enough evidence to fully understand its functioning. Although recent research is focused on “omics*”* technology (Saia-Cereda *et al.*, 2017), synaptic modulation (Shefa *et al.*, 2018) and cerebral network modelling (Faskowitz *et al.*, 2018), much of the advances on the fields of structural and functional neuroscience were made possible by the study of the cellular composition of the CNS (von Bartheld *et al.*, 2016, von Bartheld, 2018). Our comprehension of brain function and structure under both physiological normotypic (Oliveira-Pinto *et al.*, 2014; Azevedo *et al.*, 2009) and pathological conditions (Toft *et al.*, 2005; Andrade-Moraes *et al*., 2013; Repetto *et al*., 2016; Lima *et al*., 2018) has advanced pronouncedly in the last decade due to the investigation of cellular composition. Moreover, the former has been exhaustively used as the main variable of interest in some lines of research in the field of comparative neuroanatomy, allowing the development of quantitative models of brain evolution (Mota & Herculano-Houzel, 2015; Herculano-Houzel *et al.*, 2015b).

Although different methods have been developed to estimate the absolute and relative numbers of the different cell types, the stereological methodology became the most used approach. It consists of extrapolating the number of cells (events of interest) from two-dimensional histological slices of a given structure to the whole three-dimensional structure (Burke *et al.*, 2009), yielding unbiased estimates, different from earlier quantitative methods (Schmitz & Hof, 2005). Although reliable stereology demands careful experimental planning, anatomical and histological knowledge, and an expensive technical setup, being also very time-consuming, depending on the size of the analyzed structure.

In order to overcome such drawbacks, the Isotropic Fractionator (IF) method was developed (Herculano-Houzel & Lent, 2005). This technique consists of transforming structures of high architectural complexity into a homogeneous suspension of cellular nuclei, which can be stained and counted. The IF method provides estimates of the number of neurons and other cell types, such as astrocytes (Sun *et al.*, 2017), oligodendrocytes (Gomes & Guimarães *et al.*, 2018) and microglia (Chen *et al.*, 2015). It is worth noticing that this methodology was developed to process well-defined large structures that could be standardly dissected, making the IF and stereology complementary methodologies (Lent et al. 2012).

To provide evidence in support of the IF as a valid and potentially better alternative, different groups have compared the estimates obtained using the IF with the ones obtained with other techniques, such as unbiased stereology (Bahney & von Bartheld, 2014; Herculano-Houzel *et al.*, 2015a; Miller *et al.*, 2014; Ngwenya *et al.*, 2017). The overall message from these studies is that the results obtained from each method are comparable, the IF having the advantage of being faster, cheaper and having better staining with NeuN-antibodies, when it can be applied (Herculano-Houzel *et al.*, 2015a; von Bartheld, 2018), despite recent developments and guidelines for the procedure (Deniz et al., 2018).

Many studies so far employed this method to study variation in cellular composition across species, sometimes varying by many orders of magnitude. In these cases, it would require an enormous amount of methodological, random variation to weaken the main conclusions. However, in the study of smaller differences within the same species (e.g. Herculano-Houzel *et al.*, 2015b; Andrade-Moraes *et al.*, 2013; Oliveira-Pinto *et al.*, 2014), it is important to know not only if the measure has external validity – which has been repeatedly established, as just discussed above - but also whether its estimates are precise. If we are dealing with subtler differences, they could easily be masked by methodological variation that is otherwise acceptable (e.g. in cross-species studies). It could simply be that the effect size one is trying to detect is below the method’s resolution.

The amount of variation between experimenters and trials in applying the IF to the same samples has not yet been established experimentally. What concerns us here is not so much the accuracy of the measure (i.e. whether it is close to the true value being estimated), but rather the measurement error: variation across counts from the same individual and across individuals, which is important, for example, if one wants to pool together data from different sources.

Comparisons between Swiss mice and isogenic mice (C57BL/6), employing the IF method, show that estimates of neuron number for Swiss mice do seem to have more variation than estimates for isogenic mice (Figure 1). If the source of most variation was methodological noise, the variance would be the same in both kinds of mice, what suggests that the variations found in the two strains have biological origins. Additionally, a larger study on intraspecific variation in the C57BL/6 strain has been able to detect variation across individuals that amount to less than 5% of all neurons in the cerebellum (unpublished results). These observations suggest that the method is sensitive enough to detect a signal even within a species. To study the reliability of the method, we designed a set of experiments using the IF to estimate numbers of cells and numbers of neurons in mice, with different experimenters performing repeated counts using the same material, so we could address questions about sources of variation in the method.

**Figure 1.**
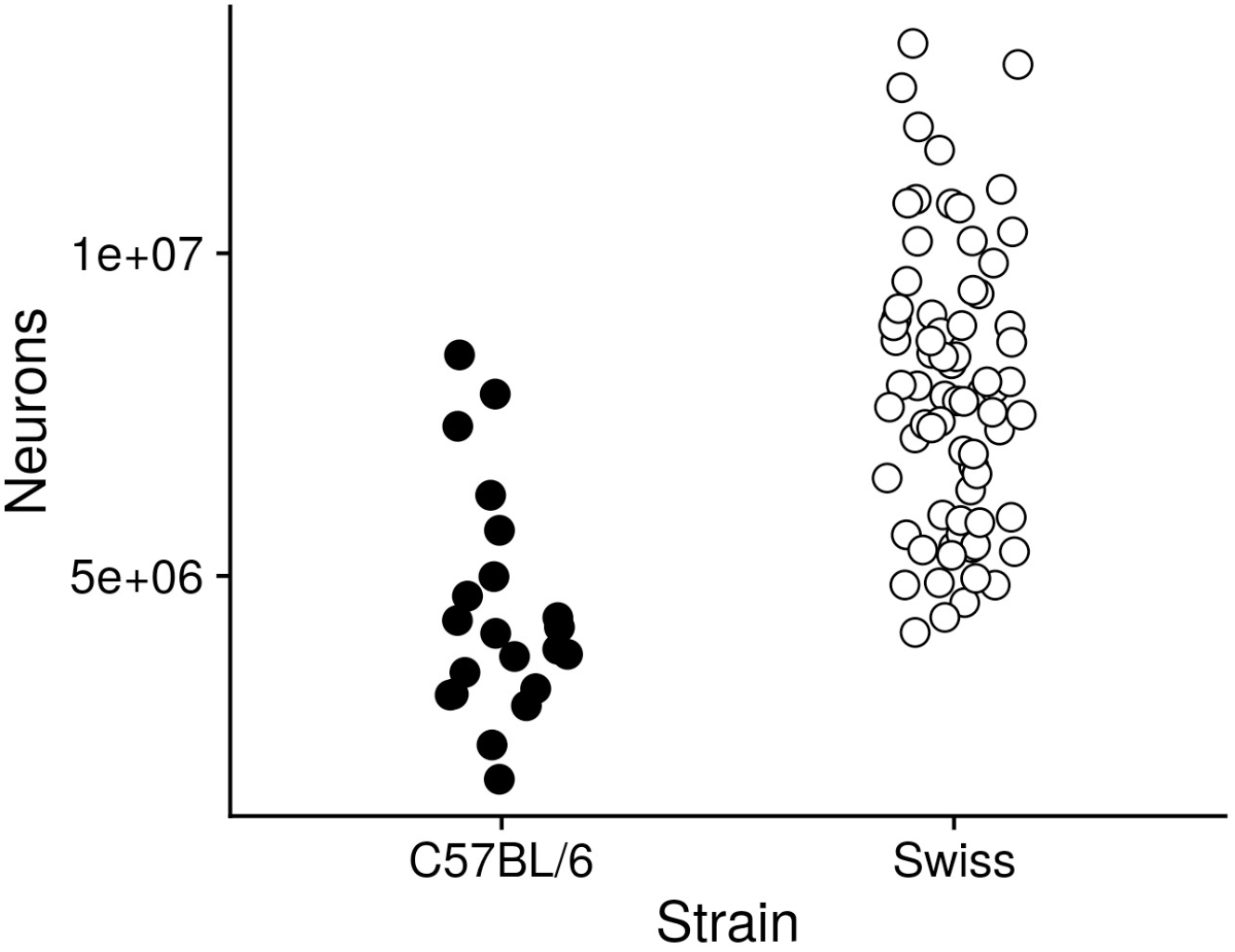
Estimates of number of cortical neurons in two strains of mice, obtained with the isotropic fractionator. Data for the Swiss strain is from Neves & Guercio et al. (2018). Data for the C57BL/6 strain is aggregated from the control group in Gomes & Guimarães et al. (2018) and from unpublished results from some of the authors. The isogenic C57BL/6 strain shows less variation in their number of neurons.

## 2 Methods

### 2.1 Experimental Design

Three Swiss mice – a non-isogenic strain, therefore having inter-individual variability – were perfused, had their brains removed from the skull and were post-fixated for two weeks in 4% paraformaldehyde (PFA) in phosphate-buffered saline (PBS). We chose to use cerebellum and cerebral cortex in this study for two reasons: (1) they are the structures with the largest cellular and neuronal densities and (2) the immunocytochemical staining of the nuclei is typically better on their neurons than elsewhere in the brain. Therefore, the cerebellum and cerebral cortex of each brain was dissected and separately dissociated through mechanical friction within a detergent solution. Once a suspension of nuclei was obtained, each of three counters – each with at least two years of experience using the isotropic fractionator – independently estimated the total number of DAPI-stained nuclei in the tissue, for each structure (cerebellum and cerebral cortex) of each of the three animals. This was done by counting samples from the tissue homogenate in a *Neubauer* chamber. Four counts from different samples of 10 μL were performed for each replicate, asL were performed for each replicate, as is usual when using the method (Herculano-Houzel & Lent, 2005). Five replicates per counter were obtained, in order to be able to assess intra-rater variability. Later, three samples of 1 mL were taken from each suspension. Each sample went through the same standard immunocytochemistry protocol for the NeuN antibody. With the samples stained with the same antibody, again, each of the three raters made repeated counts (three each) of each of the 9 samples (3 immunocytochemical stains x 3 animals) for each structure. All procedures were approved by the Ethics Committee of Animal Use in Research of Federal University of Rio de Janeiro, protocol number 046/17.

### 2.2 Isotropic Fractionator and Immunostaining

After post-fixation, the brains were processed to obtain estimates on their number of neuronal and non-neuronal cells. This is started with mechanical dissociation, to transform heterogeneous brain tissue into a suspension of cell nuclei that can be kept homogeneous by constant agitation. Nuclei can then be counted in samples from the suspension and stained by immunocytochemistry. The isotropic fractionator has been shown by at least two independent groups to give estimates comparable to stereological counts (Bahney & von Bartheld, 2014; Herculano-Houzel *et al.*, 2015a; Miller *et al.*, 2014; Ngwenya *et al.*, 2017).

Structures were mechanically dissociated in 1% Triton X-100. The resulting suspension with all nuclei was stained with the fluorescent DNA intercalant marker DAPI (4’-6-diamidino-2-phenylindole dihydrochloride, Invitrogen, USA) diluted 1:20 from a stock solution of 20 mg/L and filled with PBS 0.1M to known volume. The density of nuclei in the suspension was estimated by counting at least 4 samples of the suspension, in a *Neubauer* counting chamber, using a fluorescence microscope. The coefficient of variation between samples was typically below 0.10.

Once the estimates for cell number were obtained, a small sample of 1 mL from each suspension was incubated for NeuN, a pan neuronal nuclear marker employing the antigen Anti-NeuN Rabbit Cy3 Conjugate, ABN78C3, Millipore (Mullen *et al.*, 1992; Gittins & Harrison, 2004; RRID: AB_2314890) was incubated at 1:200 overnight with 5% Bovine Serum Albumin at 4ºC shaking, after washing thrice with 0.1M PBS. Antibodies for all the staining procedures came from the same vial. All counts, by all counters, were made in the day after the immunocytochemistry to preserve the fluorescence in equal conditions, while randomizing the order of samples and counters. Counters were blinded to which sample they were counting.

### 2.3 Data Analysis

All statistical analyses were performed in R (R Core Team, 2018). A multiple regression model was fit separately to nuclei counts and estimates of percentage of neurons, with each of these as outcomes. Predictors for the nuclei counts were animal and counter. For the percentage of neurons model, the predictors were animal, counter and also a code for the immunological staining. We performed type I analyses of variance (ANOVA) for each model. We also calculated the intra-class correlation coefficient (ICC, two-way random effects, consistency, average of raters) for each of the four procedures (cell nuclei counts and immunocytochemical staining, for cerebellum and cerebral cortex) to obtain a measure of test-retest reliability.

## 3 Results

### 3.1 DAPI counts

We start by reporting the results of the regression for DAPI counts (see Methods for details of the regression models). Differences between animals are large and consistently detectable: estimates derived from all of the three counters agree that Animal 2 has fewer cells than Animals 1 and 3, both in the cerebellum (A2-A1: −29.07, 95% CI: [−39.73, −18.41]; Figure 2A) and in the cerebral cortex (A2-A1: −9.52, 95% CI: [−15.32, −3.72]; Figure 2B). Regarding differences between counters, the regression shows that counters may have biases larger than the differences between animals. In our small sample, Counter 1 counts fewer cells for every animal, compared to the other counters, which have much more similar counts. For instance, the difference between Counters 1 and 3 for the cerebral cortex is 18.17 (95% CI: [12.37, 23.97]) and for the cerebellum, 11.42 (95% CI: [0.76, 22.08]). The estimated differences between Counters are larger than the largest difference between animals (see Table 1 for details).

**Table 1.** Regression summary for DAPI counts.

| **Cerebellum** |  |  |
| --- | --- | --- |
|  | **Estimate** | **Std. Error** |
| Intercept | 255.138 | 4.965 |
| Animal 2 | -29.067 | 5.439 |
| Animal 3 | 5.000 | 5.439 |
| Counter 2 | 14.667 | 5.439 |
| Counter 3 | 11.417 | 5.439 |
| **Cerebral Cortex** |  |  |
|  | **Estimate** | **Std. Error** |
| Intercept | 109.800 | 2.701 |
| Animal 2 | -9.516 | 2.959 |
| Animal 3 | -1.283 | 2.959 |
| Counter 2 | 23.733 | 2.959 |
| Counter 3 | 18.167 | 2.959 |

**Figure 2.**
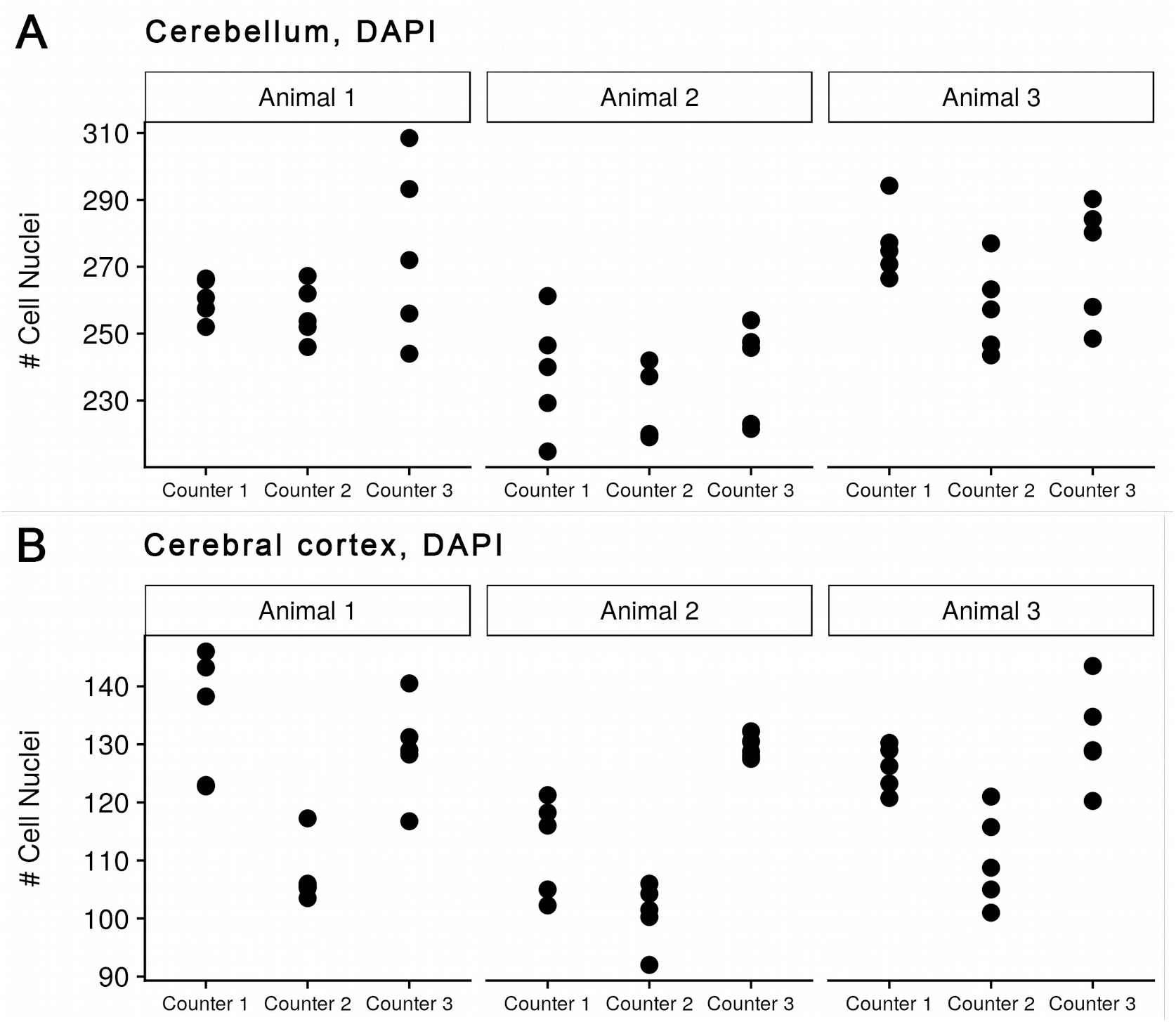
Counts of DAPI-stained cell nuclei for each combination of animal and counter, for the cerebellum (A) and cerebral cortex (B). N = 5 replicates for each combination. While samples from Animal 2 are consistently found to have less cells by different counters, systematic differences between counters and variation within replicates are not negligible compared to the difference between animals.

An analysis of variance (ANOVA) shows that for the cerebral cortex most of the variance is at the Counter level (η = 0.58), however, the percentage of variance within animals (η = 0.1) is much smaller than residual variance (η = 0.33), suggesting that high intra-rater variability could be of concern. While coefficients of variation (CV) are low (typically below 10%), in absolute numbers, the standard deviations of the intra-rater counts range from very small to as large as the differences in counts between animals (e.g., for the cerebral cortex, the 5 replicates for Counter 3 on Animal 1 have a mean of 134.66 with a standard deviation of 11.10). For the cerebellum, the ANOVA shows that, on the contrary, most of the variation is at the Animal level (η = 0.49), not at the Counters level (η = 0.09), although the difference between counters is similar to the one found for the cerebral cortex. This is due to the larger absolute differences between animals for the cerebellum. Nevertheless, residual variance is still considerably large (η = 0.43).

We also calculated the test-retest reliability of the repeated cell counts by the same counters, which were found to be high. For the cerebral cortex, the coefficient is 0.93 (95% CI: [0.81, 0.98]) and for the cerebellum, it is 0.85 (95% CI: [0.62, 0.96]).

### 3.2 Immunocytochemical staining counts

For the counts of immunocytochemically stained nuclei, there are two differences compared to the DAPI analysis: (1) outcomes are expressed as percentages, from the ratios of the number of NeuN-positive nuclei to the number of DAPI stained nuclei and (2) there is an extra modeled source of variation, namely the immunocytochemical staining procedure, which is much more capricious than DNA intercalation by DAPI.

Estimates from the model suggest that systematic differences between animals and counters are small – in fact, standard errors on the estimates are of same magnitude of the estimates themselves, sometimes larger (see Figure 3 and Table 2). Moreover, most of the variance is left unexplained by the model (residual variance is 0.80 for the cerebral cortex and 0.82 for the cerebellum). Standard deviations within replicate counts from the same stained sample and counter range from 0.69 to 5.64 in the cerebral cortex and from 0.69 to 12.95. The overall picture is that variation in the immunological staining counts is random, not systematic. Within our small sample, this random variation is larger than the variation between animals. No biases among counters or large differences between animals or between staining intensity were found.

**Table 2.** Regression summary for immunological staining counts.

| **Cerebellum** |  |  |
| --- | --- | --- |
|  | **Estimate** | **Std. Error** |
| Intercept | 67.394 | 2.059 |
| Animal 2 | 1.677 | 2.633 |
| Animal 3 | 0.096 | 2.633 |
| Counter 2 | 1.120 | 1.520 |
| Counter 3 | 0.540 | 1.520 |
| Animal 1: Imuno 2 | 2.482 | 2.633 |
| Animal 2: Imuno 2 | -2.056 | 2.633 |
| Animal 3: Imuno 2 | -5.861 | 2.633 |
| Animal 1: Imuno 3 | 0.217 | 2.633 |
| Animal 2: Imuno 3 | 1.476 | 2.633 |
| Animal 3: Imuno 3 | -0.432 | 2.633 |
| **Cerebral Cortex** |  |  |
|  | **Estimate** | **Std. Error** |
| Intercept | 26.952 | 1.370 |
| Animal 2 | -0.820 | 1.752 |
| Animal 3 | -2.973 | 1.752 |
| Counter 2 | 1.604 | 1.012 |
| Counter 3 | 2.477 | 1.012 |
| Animal 1: Imuno 2 | -1.957 | 1.752 |
| Animal 2: Imuno 2 | 1.444 | 1.752 |
| Animal 3: Imuno 2 | 2.746 | 1.752 |
| Animal 1: Imuno 3 | -2.676 | 1.752 |
| Animal 2: Imuno 3 | -1.376 | 1.752 |
| Animal 3: Imuno 3 | -0.959 | 1.752 |

**Figure 3.**
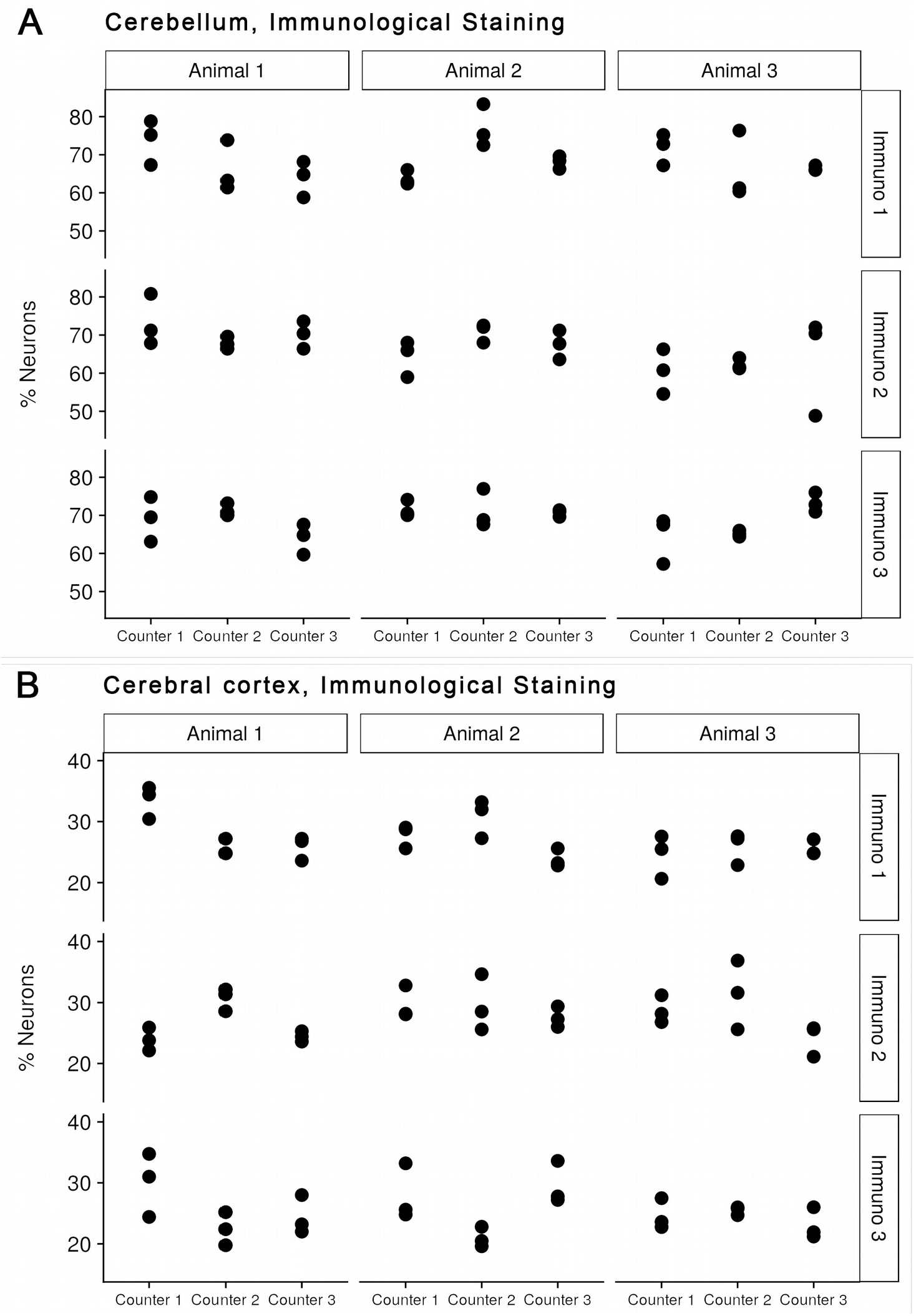
Counts of the NeuN-stained cell nuclei for each combination of animal, counter and immunostaining, for the cerebellum (A) and cerebral cortex (B). N = 3 replicates for each combination. Most of the variation in these counts appear not to be systematic.

We calculated the test-retest reliability for the immunocytochemical staining counts as well. For the cerebral cortex, we found a coefficient of 0.71 (95% CI: [0.45, 0.86]) and for the cerebellum, of 0.51 (95% CI: [0.09, 0.76]). These are lower than the coefficients for the DAPI counts. However, the reliability for the cerebral cortex is still considered high. For the cerebellum, the test-retest reliability is possibly much lower, even if the quality of the immunocytochemical staining be usually better, which should reduce ambiguity in counts.

## 4 Discussion

We sought to identify sources of error in the estimates obtained with the isotropic fractionator method for counting neurons and total cells in brain tissue. For total cell counts, we found large systematic differences between counters. Nevertheless, an estimated difference of around 10% (amounting to 30 nuclei counted in cerebellum samples and 9 in cerebral cortex samples) between the animals was consistently detectable by all three counters. For the neuronal cell counts, we found that the extra source of variation – the immunocytochemical staining – is random and comparable to the estimated differences between animals and counters.

Although the present study is limited in scope with a small sample size, the data suggest a few considerations to be kept in mind when using the method, in order to reduce error in the measurements. First, we found large, systematic biases between counters. For individual projects this is unlikely to be a problem: the same person might be responsible for all counts – this was the case for most of the authors’ past projects using the method – or in cases of more than one counter, they can calibrate their counts until they consistently agree. This can be a problem, nevertheless, when aggregating and comparing data from the published literature, since the estimates were obtained by different counters, in different research groups. In this case, these systematic biases between counters are a source of error we must account for, especially if one is studying small differences. Using flow cytometry to obtain the counts could be one way around this issue. Although it has been used in some studies with the technique (e.g. Young et al., 2012), costs are a major difficulty and perhaps may not significantly ameliorate for a second global issue: the reliability of immunocytochemical markers. In addition, flow cytometry counters do not discern between healthy nuclei and nuclear fragments, what can be a source of error avoided by human counters. The second consideration regards the random error coming from the immunocytochemical counts. It is unlikely that the counting differences are caused by biases from the counters because the variation does not seem to be systematic between them. On the other hand, antibodies are a known source of error in the biomedical sciences (Baker, 2015). In this study, all staining procedures were conducted simultaneously, following the same protocol, with all the antibody used coming from a single vial. If is the case that the random variation observed comes from differences in the staining effectivity, we imagine that the error can be even larger, again, when comparing neuronal counts obtained with antibodies in less controlled conditions – reagents stored differently, from different brands, with different fluorophores, different staining protocols, etc. Notice, though, that while noisy, the estimates of the percentage of neurons in the cerebellum are consistently higher than the percentages found for the cerebral cortex, agreeing with the literature (Herculano-Houzel, 2010). Small differences in the percentage of neurons, though, could be unreliably detected by this method. In this scenario, one might get better estimates from total cell counts only, since the counts from immunological staining might be adding random error comparable or larger than the effect size of interest to detect.

If differences are large – such as in comparisons between different species (Herculano-Houzel *et al.*, 2014) –, these concerns are less important. Still, we gain by counting large numbers of nuclei. If the counting error is independent of number of cells counted (or weakly correlated with it), by counting a larger number of nuclei, errors become less consequential. For instance, for this study, every structure had its nuclei homogenized into a volume of 14 mL. At this density, the difference between animals was about 10 nuclei counted in the *Neubauer* chamber (out of 100-150 nuclei counted in total, per chamber). If we had halved the volume, the density would double and we would count 200-300 nuclei per chamber, so the difference between animals would become of 20 nuclei. Then, assuming the error is still the same, it would pay to count samples with a large number of nuclei, since the error would be smaller compared to the expected effect.

All of this speaks to the importance of planning the experimental design beforehand, including the statistical analysis to be performed. To use this method – or any method, for that matter – it is fundamental to have a grasp of the expected effect size in comparison with the method’s precision, obtained from *a priori* reasoning and independent data (Gelman & Carlin, 2014). This will improve the odds of obtaining a good estimate – i.e. an estimate where the error is not as large as the estimate itself.

The main limitation of the present work is that all the experimenters had previous experience with the methods, each one having at least 5 years. Due to the subjective non-automated nature of the counts, it is expected higher variance between unexperienced counters. Nevertheless, it is feasible to learn the method by oneself, if trained by a senior counter (by counting and recounting the same microscope fields). In our experience, a month of regular and frequent training is enough to produce consistent data. Still, another experimental design, including naïve counters would be of great value, as it would allow to gauge not only the difference of rating between experienced and naïve raters but also consistence among naïve experimenters.

Investigating the reliability of our own methods is one of the defining characteristics of science (Ioannidis, 2012). Methodological variability is understudied in the biomedical sciences and the attempts that have been made to systematically investigate it have shown that there are large amounts of methodological variations among labs (Crabbe, 1999; Hines *et al.*, 2014). We agree with von Bartheld (2018, page 10), that “more work is needed to identify and to minimize sources of bias” in counting methods. This study puts the isotropic fractionator among the methods for which there is data regarding their reliability. We regard this type of methodological studies as greatly beneficial for science in general, since for many commonly used methods there is no systematic characterization of their precision, which means we might be using them broadly to investigate effect sizes they could not possibly detect (Loken & Gelman, 2017). It is imperative that we better understand the reproducibility of our methods and improve our reporting of how procedures are performed (von Bartheld, 2018). Hopefully, the many reproducibility efforts started in recent years will fill this gap (Errington *et al.*, 2014; Amaral *et al.*, 2019). Although limited, this study is a first step in the right direction for this cell counting method, especially in this moment, as biomedical science goes through a credibility revolution (Munafò *et al.*, 2017).

## Author Contributions

K.N., D.M., D.R., B.V-G., B.M., R.L. designed the research. K.N., D.M., D.R., B.V-G. and P.M.I. perfomed experiments, analysed data and wrote the manuscript. B.M. and R.L. supervised the project and revised the manuscript.

## Abbreviations

CNS: Central Nervous System
IF: Isotropic Fractionator
PFA: Paraformaldehyde
PBS: Phosphate Buffered Saline
DAFI: 4’-6-diamidino-2-phenylindole dihydrochloride
BSA: Bovine Serum Albumine
ICC: Intra-class correlation coefficient
CV: coeficiente of variation
DNA: Deoxyribonucleic acid

